# Systematic Analysis of Human Colorectal Cancer scRNA-seq Revealed Limited Pro-tumoral IL-17 Production Potential in Gamma Delta T Cells

**DOI:** 10.1101/2024.07.18.604156

**Authors:** Ran Ran, Martin Trapecar, Douglas K. Brubaker

## Abstract

Gamma delta (γδ) T cells play a crucial role in anti-tumor immunity due to their cytotoxic properties. However, the role and extent of γδ T cells in production of pro-tumorigenic interleukin-17 (IL-17) within the tumor microenvironment (TME) of colorectal cancer (CRC) remains controversial. In this study, we re-analyzed nine published human CRC whole-tissue single-cell RNA sequencing (scRNA-seq) datasets, identifying 18,483 γδ T cells out of 951,785 total cells, in the neoplastic or adjacent normal tissue of 165 human CRC patients. Our results confirm that tumor-infiltrating γδ T cells exhibit high cytotoxicity-related transcription in both tumor and adjacent normal tissues, but critically, none of the γδ T cell clusters showed IL-17 production potential. We also identified various γδ T cell subsets, including Teff, TRM, Tpex, and Tex, and noted an increased expression of cytotoxic molecules in tumor-infiltrating γδ T cells compared to their normal area counterparts. Our work demonstrates that γδ T cells in CRC primarily function as cytotoxic effector cells rather than IL-17 producers, mitigating the concerns about their potential pro-tumorigenic roles in CRC, highlighting the importance of accurately characterizing these cells for cancer immunotherapy research and the unneglectable cross-species discrepancy between the mouse and human immune system in the study of cancer immunology.

## Introduction

### γδ T Cells in Colorectal Cancer: An Introduction

T lymphocytes are pivotal to adaptive immunity, specializing in initiating targeted immune responses against diverse antigens. Most T cells in the human body are alpha beta T cells (αβ T cells), whose T-cell receptors (TCRs) are composed of an α chain and a β chain. These cells rely on the major histocompatibility complex (MHC) in other cells to present antigens for TCR binding and subsequent activation^1^. Gamma delta T cells (γδ T cells) are a less common population with a TCR composed of γ and δ chains^2^. γδ T cells typically account for less than 5% of T lymphocytes in the blood, but this proportion is higher in subcutaneous tissue^3^, as well as the mucosa of the intestinal^4^, respiratory^5^, and urogenital tracts^6^.

Activation of γδ T cells can occur through a TCR-dependent process like that of their αβ counterparts but without MHC-mediated antigen presentation. The Vδ2 subset recognizes phosphoantigens modified by the BTN2A1-BTN3A1 complex^7^. Non-Vδ2 TCRs recognize CD1 family members and MR1, and are not necisarily antigen dependent^8^. Natural killer (NK) receptors including NKG2D, NKp30, and DNAM-1 on the γδ T cells surface recognize MICA/B, ULBP, B7-H6, and Nectin-like-5^9^, allowing them to be activated in a TCR-independent, innate-like manner^10^. These unique antigen recognition properties endow γδ T cells with significant potential to maintain homeostasis^11^ and execute anti-tumor immune responses^12,13^. They have been observed to infiltrate various tumor types, including rectal^14^, breast^15^, pancreatic^16^, kidney^17^, and colorectal cancers (CRC)^18,19^, exhibiting anti-tumoral cytotoxicity and regulatory effects in co-culture experiments *in vitro*^20–29^, *in vivo* animal models^30–39^, and functional assays on patient-derived γδ T cells^40,41^. γδ T cells can recognize and lyse cancer cells, which often exhibit MHC deficiencies or abnormalities^42^, in an MHC-unrestricted manner by producing classical cytotoxic molecules like granzyme B and perforin through direct contact via death receptor signaling^25,43,44^.

CRC is the third most frequently diagnosed cancer and the second leading cause of cancer deaths^45–47^. It is confined to the colon or rectum and is marked by the abnormal growth of glandular epithelial cells^48^. Surgery is the primary treatment strategy for advanced CRC, but 71.2% of advanced CRC patients recur within two years after surgery, and the five-year survival rate is only 34.7%^49,50^. The invention of chimeric antigen receptor-T (CAR-T) cells dramatically shifts the paradigm of cancer therapy, especially malignancies like leukemia and melanoma^51^. However, their efficacy in solid tumors like CRC is limited by poor tumor trafficking and infiltration, the presence of an immunosuppressive TME, and adverse events associated with such therapy^51,52^. Given the association of γδ T cells with antitumor activity in various cancers, including CRC, which occurs in the colon—one of the most abundant sites of γδ T cell residence—γδ T cells have been explored and engineered as an anti-tumor strategy against CRC. Current engineered γδ T cell therapies exhibit significantly enhanced potential to achieve a targeted cell elimination and produce cytokines, leading to more substantial decreases in tumor size and inhibition of tumor progression compared to non-engineered γδ T cells^53–56^.

### The Controversy Around γδ T Cells’ Pro-Tumoral Effect on Human Colorectal Cancer

There is debate over the potential pro-tumoral activities of human γδ T cells motivated by their apparent production of interleukin-17 (IL-17). IL-17 can promote epithelial-mesenchymal transition, enhance tumor survival, and attract myeloid-derived suppressor cells (MDSCs) to establish an immunosuppressive environment conducive to tumor growth^57^. In 2014, the first evidence emerged purporting that tumor-infiltrating γδ T cells producing IL-17 (hereafter: γδ T17), but not T helper17 (Th17) or cytotoxic T cells producing IL-17 (Tc17), are the major IL-17A (hereafter: IL-17)-producing cells in human CRC^58^. Wu et al. used flow cytometry to show that the percentage of IL-17A^+^CD3^+^ cells in the CD45^+^ population increased from 1.48% in normal tissue to 6.98% in tumor tissue. Gating of the CD3^+^ T cells into CD8, CD4, and TCRγδ populations revealed a higher proportion of IL-17A^+^ cells among TCRγδ^+^ T cells compared to CD8^+^ and CD4^+^ T cells. Wu et al. also showed that cytokines from γδ T17 cells enhance the attraction, growth, and survival of MDSCs using transwell co-culture assays, reporting less T cell proliferation, less IFN-γ production, and increased MDSC migration. In 2017, the same group published another paper on tumor-infiltrating CD39^+^ γδ Tregs in human CRC^59^ and reported IL-17 production in CD39^+^ γδ Tregs. In 2022, following these lines of evidence, Reis et al. performed scRNA-seq paired with γδ scTCR-seq on sorted anti-TCRγδ^+^CD3^+^ T cells isolated from human CRC patients^60^. Reis et al. found 2 tumor-infiltrating cell clusters in humans enriched with CD9 and LGALS3, genes identified by others as markers of murine IL-17 producing γδ T^61^. Their subsequent experiments in mice showed that it is mainly the PD-1^+^ γδ T cells that produced IL-17.

However, there is evidence that contradicts the assertion that γδ T17 cells exist in high abundance in human CRC tissue. Although the presence of γδ T17 is supported by strong evidence in murine CRC models^13,62–69^ and reviewed in detail elsewhere^70,71^, there is growing consensus that, in contrast to mice, γδ T17 cells are not present in significant numbers in the healthy human colon^72–75^. While γδ T17 cells have been observed in other human diseases in very small amounts^76–79^, studies on CRC suggest their presence is statistically insignificant. For instance, Meraviglia et al. evaluated the CRC-infiltrating γδ T cells’ ability to secrete IL-17, IFN-γ, and TNF-α upon the stimulation of ionomycin and PMA *in vitro* using different FACS gating strategies^18^. The result showed that most CD45^+^ IL-17^+^ cells in both CRC and adjacent normal tissues were CD3^+^ αβ T cells while the γδ T cells were producing IFN-γ. Additionally, Amicarella et al. performed IL-17 staining by immunohistochemistry on 1,148 CRC tumors and used flow cytometry to examine the phenotypes of IL-17-producing cells, and they found less than 1% of these cell types showed evidence of IL-17 production^19^.

The consequences of this controversy over the presence of pro-tumor, IL-17 producing γδ T cells in CRC are far-reaching and have implications for the development of γδ T cell-targeting CRC therapies. Based on the current evidence, we hypothesized that γδ T17 cells do not represent a meaningful, naturally occurring population of γδ T cells in human CRC. Motivated by the approach of previous studies that characterized the IL-17-producing behavior of γδ T cells using scRNA-seq, we sought to determine whether IL-17 transcription can be identified in γδ T cells by analyzing and integrating whole tissue human CRC data across nine published studies. Our approach involved a detailed examination of the cell clusters in Reis et al.’s study that expressed genes related to IL-17-producing γδ T cells. Subsequently, we analyzed scRNA-seq datasets from flow cytometry-sorted, purified lymphoid cell populations to establish reliable transcriptomic markers for γδ T cells. Finally, we analyzed multiple published human CRC scRNA-seq datasets to computationally isolate γδ T cells and assess their potential to produce IL-17 (Figure 1A).

**Figure 1:**
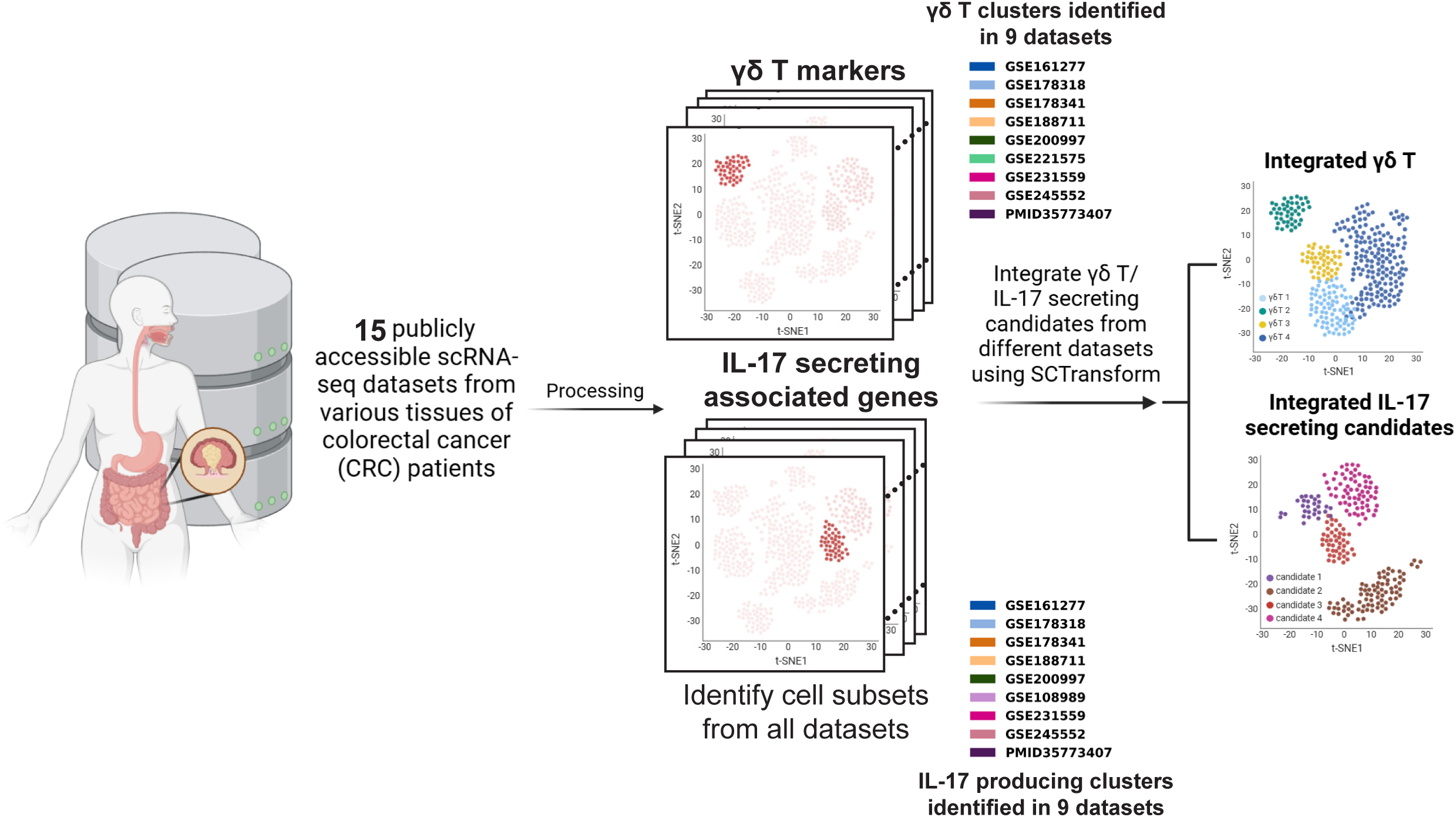
Overview of the Data Collection and Integration. Schematic of data collection, processing, cell selection, and integration.

## Results

### Previously Identified Human IL-17–Producing γδ T Cell-like Cells Identified Lack Expression of Genes Characteristic of IL-17 Production

In their work, Reis et al. stated that “…tumor-infiltrating γδ T cells shown an overall increased cytokine signature (gene ontology: 0005126), including IL-17–producing γδ T cell-related genes, such as enriched in clusters 2 and 5, when compared with cells isolated from adjacent nontumor areas^60^.” We obtained the unaltered expression matrix and cluster annotation from Reis et al.’s original paper and queried the IL-17-producing related genes in each of the clusters. Specifically, we examined IL-17 production from three perspectives: (1) the detectable transcripts of the IL-17 cytokine family, (2) the expression of members in the transcriptional complex of IL-17^80^, and (3) previously identified features used as indicators of IL-17 production potential, or the gene ontology for the calculation of an “IL-17 score” used by Sanchez Sanchez et al.^81^ and Tan et al^61^.

Surprisingly, clusters 2 and 5 exhibited very low levels of IL17A, IL17F, master transcription factor RORC, and other genes important to IL-17 transcription, such as IL23R^82^, CCR6^83^, and MAF^84^ (Figure S1A-B). While IL-17 expression could be hard to capture in the scRNA-seq data, depending on the sensitivity of the platform, RORC transcripts were prevalently captured in other studies that generated scRNA-seq data in known IL-17-producing cells like T helper 17 (Th17), Mucosal-associated invariant T (MAIT) cells, and innate lymphoid cell (ILC)^85,86^. Given that RORγt, encoded by RORC, is needed for the transcription of IL-17^76^, the missing RORC is likely not due to the technical limitation of scRNA-seq, raising a question about IL-17 production in clusters 2 and 5. Indeed, the annotation of IL-17–producing γδ T-like cells by Reis et. al is based on the expression of two genes, CD9 and LGALS3, two features of a murine γδ17 subset from previous studies.

In cluster 4, marginal IL-17A and RORC expression was noted (Figure S1A-C). However, these cells also expressed helper T cell features like CD40LG and CXCR5, regulatory features like CTLA4 and IL2, immune activation markers such as OX40 and TNF, and a low level of CD4, features characteristic of TCRαβ Tfh/Treg cells. Notably, these cells did not express the classical γδ T marker TRDC, showed almost no productive TCRγδ chains in the scTCR-seq, and had marginal TRBV5-1 and TRBV20-1 expression, likely indicating TCRαβ identity (Figure S1C). Moreover, despite using a seemingly reasonable CD3^+^TCRγδ^+^TCRαβ^-^ gating strategy, Reis et al. found abundant TRBV6-1 and TRBV21-1 transcripts in TRDC^-^ clusters 7 and 8. This results together with dropout of TCRγδ-seq in these cluster raises questions of the purity of γδ T cells in their study.

### Creating a Reliable Gene Set for Classification of γδ T in scRNA-seq

After we examined Reis et al.’s data, we asked how γδ T cells in other scRNA-seq studies behave. To accurately identify them in the CRC microenvironment, we needed to establish reliable γδ T transcriptomic markers. Typically, TRGC1/2, TRDC, and TRDV1/2/3 are used as marker genes for γδ T cells, both in studies with manual cell type annotation and with auto-annotators like CellTypist^87–91^. Recently, Zheng Song et al. developed a TCR module scoring strategy based on all mappable constant and variable αβ and γδ TCR genes^92^. However, given the plasticity of T cells, the CRC γδ T phenotype can be dramatically influenced by the complex environmental cues in the TME.

Instead of using an inclusive list of γδ T surface molecules, secreted cytokines, and differentially expressed genes from individual studies—which may not be fully captured in a single sequencing run or may be partially inactive in the TME considering the limited knowledge about CRC γδ T cells—we developed a concise set of core features specific to CD3 and the γδ TCR. Our criteria included a critical review of the existing literature, re-analysis of scRNA-seq data from tumors which had been flow cytometry-sorted before sequencing, and reanalysis of tumor data with paired with scTCR-seq, allowing us to combine multiple orthogonal lines of experimental evidence to establish our new set of reliable γδ T cell markers. Due to the limited access to human tissues and the cost of such experiment procedures, datasets that meet such requirements and are publicly available are not common. One of them is provided by Gheradin et al., who conducted scRNA-seq with paired scTCR-seq (both αβ and γδ) on sorted anti-CD3^+^ T cells isolated from human Merkel cell carcinoma^17^, and another is provided by Rancan et al., who performed scRNA-seq paired with TCRαβ-only scTCR-seq on the anti-CD3^+^ T cells isolated from human kidney cancer^93^.

TRDC, the T Cell Receptor Delta Constant region, has long been used as a γδ T marker in scRNA-seq analysis. Even for cells that have passed the gating of CD3^+^TCRγδ^+^ in pre-sequencing flow cytometry but are TRDC^-^ in the scRNA-seq count matrix, it is common practice to exclude them from downstream analysis^17,91^. Cells from Gheradin et al that do not express TRDC also have no TRD chain captured (Figure S2A). The finding of productive TRA/TRB chains in these TRDC^-^ cells further supports the idea that they are more likely αβ T cells (Figure S2A). Similarly, in Rancan et al.’s kidney cancer T cell data, the loss of TRDC highly correlates with the detection of productive TRA/B chains (Figure S2B). Therefore, we posit that TRDC is a canonical transcriptional marker for γδ T cells, and T cells without TRDC expression are less likely to be actual γδ T cells and cannot be reliably classified as γδ T cells by RNA expression alone.

The CD3 complex transmits the activation signal into the T cell. The high and consistent expression of all four CD3 subunits—CD3D, CD3E, CD3G, and CD247 (CD3Z)—in cell clusters distinguishes γδ T cells from their lymphoid relatives like NK and ILCs, which share similar early developmental trajectories and cellular programs with γδ T cells^94,95^. Therefore, it is not surprising to see NK and ILCs also highly express TRDC RNA^88^ and occasionally show expression of cytoplasmic CD3s^96,97^. However, scRNA-seq analysis of CD3^+^ T cells showed they are usually CD3D/E/G/Z quadruple positive, while NK/ILC only express parts of them at low levels (Figure S2C). Thus, the co-expression of all four CD3s is a strong indicator of T cell identity, which aligns with flow cytometry results^96^. Taken together, we conclude that the combination of CD3D, CD3E, CD3G, CD247, and TRDC constitute a reliable, minimal set of marker genes to distinguish γδ T cells from unsorted whole tissue scRNA-seq.

### γδ T Cells in Human Colorectal Cancer Do Not Have Phenotypic Characteristics of IL-17 Production

Having established a reliable set of markers for classifying γδ T cells in human CRCs, we developed a data integration workflow to assess the presence and function of these cells in a comprehensive survey of the human CRC scRNA-seq dataset (Figure 1A). We searched for publicly available scRNA-seq datasets from treatment naïve CRCs in indexed journals and Gene Expression Omnibus, and we obtained 15 published human CRC whole-tissue scRNA-seq datasets collected from patients of varying genders, ages, tumor stages, and tumor locations for analysis.

In each study, cell clusters that highly co-expressed CD3D, CD3E, CD3G, CD247, and TRDC were identified as γδ T cells. Distinct γδ T cell clusters with more than 50 cells were identified from 9 studies that provided raw counts. A total of 18,483 γδ T cells were identified from the neoplastic and adjacent normal tissues of 165 human CRC patients and integrated (Figure S3A). The integrated data includes γδ T cells sampled from a diverse range of age, sex, and sampling sites, providing insight into their common behavior in human CRC (Figure S3A). Note that GSE161277 sampled both carcinoma and adenoma tissue, which allowed us to assess IL-17 production in adenoma as well as carcinoma samples.

As we checked the gene expression of 3 sets of IL-17-producing features in Reis et al.’s clusters, we also examined their expression in the integrated γδ T cells (Figure 2). First, none of the identified TRDC^+^ γδ T cells exhibited significant (>1% of the cluster) IL-17A-F expression (Figure 2C). We next assessed expression of the IL-17 master transcription factor RORC^98–101^ and other components of the transcriptional complex, such as RUNX1, IRF4, BATF, KLF4, and STAT3(Figure 2C)^80^. RORC, RUNX1, and KLF4 were undetectable in all γδ T cells. IRF4 was marginally expressed in exhausted T cells (Tex), and BATF was moderately expressed in non-memory-like populations. STAT3 was universally expressed in all γδ T cells. Although there are activities of IRF4, BATF, and STAT3, these transcription factors are involved in multiple signaling pathways and are not specific to IL-17 production^102–104^. The absence of RORC—which is required for the IL-17 production^76^, highly expressed in IL-17-producing human T cells, and commonly detectable by scRNA-seq^85,86^—in all γδ T cells across all 9 studies suggests that RORC is truly not expressed and that this result is not due to technical dropouts or temporary transcriptional quiescence. Finally, we queried the expression of IL-17 secretion-related features from other groups, summarized by Sanchez Sanchez et al.^81^ and Tan et al.^61^, including IL23R^82^, CCR6^83^, MAF^84^, and AHR^105^, which are strong indicators of a cell’s IL-17-producing identity. These markers were also all absent in the γδ T cells in our combined dataset (Figure 2C).

**Figure 2:**
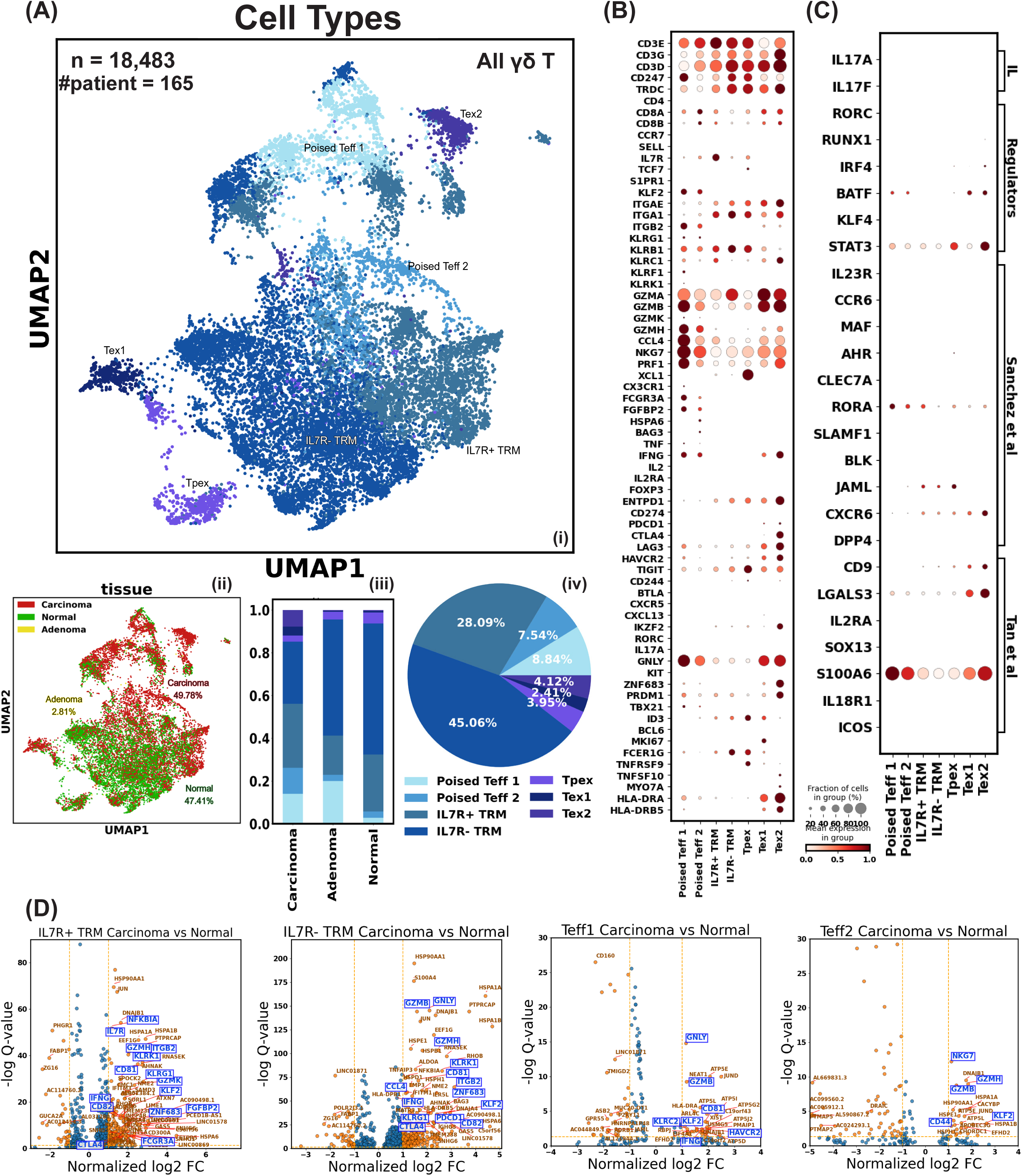
The Landscape of Human γδ T Cells in Colorectal Cancer. A) i) Uniform manifold approximation and projection (UMAP) of γδ T cells showing subsets: Teff (effector-like T cells), TRM (tissue-resident memory T cells), Tpex (progenitor exhausted T cells), and Tex (exhausted T cells). ii) UMAP of γδ T cells colored by tissue origin. iii) Composition of γδ T cell subtypes identified from different tissue types. iv) Composition of γδ T cell subtypes in the integrated dataset. B) Dot plot displaying the level and percentage of each γδ T cell subtype expressing key T cell features related to identity, mobility, tissue residency, cytotoxicity, and exhaustion. C) Dot plot showing the level and percentage of each γδ T cell subtype expressing IL-17 production-related features. IL: interleukin, direct IL-17 transcript; Regulators: proteins regulating IL-17 transcription; Sanchez et al.: IL-17 producing features as defined by Sanchez Sanchez et al.; Tan et al.: IL-17 producing features from Tan et al.’s mouse study, with CD9 and LGALS3 signatures used by Reis et al. to annotate their data. D) Volcano plot of differentially expressed genes. X-axis: gene expression level log2 fold change (log2 FC) in γδ T cells found in carcinoma tissue versus adjacent normal tissue. Y-axis: -log10 q-value (false discovery rate) of gene’s fold change in γδ T cells in carcinoma tissue versus normal tissue. The log2 FC threshold for a gene to be considered differentially expressed was set at 1.

### Human Colorectal Cancer-Related γδ T Cells Are Heterogeneous

Having generated a comprehensive dataset of γδ T cells subjected to scRNA-seq and found no evidence for IL-17 production, we leveraged our dataset to classify the subtypes and potential functions (Figure 2A). We identified four distinct subtypes of γδ T cells in our integrated dataset: poised effector-like T cells (Teff), tissue-resident memory T cells (TRM), progenitor exhausted T cells (Tpex), and Tex.

Teff cells exhibited classical signs of effector function, such as differentiation marker KLRG1, FcγRIIIa receptor FCGR3A (common in macrophages, NK cells, and γδ T cells), fibroblast growth factor binding protein 2 FGFBP2 (related to cytotoxicity), and transcription of classical and well-studied molecules TNF, IFNG, and GZMK. They also expressed the transcription factor T-bet, which guides effector differentiation, and had low IL7R (CD127), indicative of a limited lifetime (Figure 2B). The limited expression of integrin ITGAE and ITGA1 suggested a reduced degree of anchoring to neighboring cells, while the absence of selectin-L (SELL) and CCR7 hinted that these cells were not likely to enter circulation and home back to the lymph nodes, indicating they were likely poised in the tumor. Indeed, the genes expressed by these poised cells greatly resembles the previously reported CD69^+^ITGB2^+^ TRM, bona fide TRM cells that are also IL7R^-^ and ITGAE^-106^

The poised Teff cells could be further divided into a more effector-like population (Teff 1) and a more quiescent, TRM-like population (Teff 2) (Figure 2B). Teff 1 uniquely transcribes KLRF1, related to NK cell cytotoxicity, KLRK1 (NKG2D), mediating TCR-independent activation in γδ T cells, and CX3CR1, indicating differentiation during the effector phase and robust cytotoxicity in antiviral immunity. Teff 2 shows higher ITGA1 and marginal levels of ID3, implying the start of a tissue residency program. It uniquely expressed heat shock proteins HSPA6 and HSPA1A, and stress response genes DNAJA4 and DNAJB1^107,108^.

The TRM population is characterized by high expression of ITGAE and ITGA1 and low transcription of CCR7, SELL, S1PR1, and KLF2, implying limited circulation ability (Figure 2B). We further divided TRM based on differential IL7R expression following previous literature^109^. A higher portion of TRM found in carcinoma tissue is IL7R^+^, while it is mainly IL7R^-^ TRM in normal and adenoma tissue. IL7R^+^ TRM has higher KLRC1, encoding the NK cell inhibitory receptor NKG2A, which binds HLA-E and transmits inhibitory signals that impair NK cell function^110^. The IL7R^-^ TRM has higher FCER1G, marking innate-like αβ T cells with cytotoxic and tumor-infiltrating potential^111^. Both TRM populations showed low transcription of TNF and IFNG.

Tpex cells, which emerge during chronic infections or in tumor environments, are a subset of memory-like T cells exhibiting the hallmarks of Tex cells^112,113^. They retain the ability to proliferate and differentiate into Tex cells under repetitive antigen stimulation, sustaining a reduced but persistent immune response against antigens that the body has failed to clear^112,114^. Tpex cells are marked by partial expression of exhaustion markers such as HAVCR2 (TIM-3),

LAG3, TIGIT, and CD244 (2B4) (Figure 2B). Importantly, they express TCF7 (TCF-1), highlighting their differentiation potential. The moderate expression of IL7R adds to their memory-like longevity. They also uniquely and highly express XCL1, the ligand of XCR1, a receptor confined to myeloid dendritic cells (DC), suggesting enhanced interaction with DCs. This aligns with previous descriptions of their function^115^. Additionally, Tpex cells display high tissue residency markers ITGAE/ITGA1, FCER1G (correlating with tumor infiltration and effector function), and T cell activation markers TNFRSF9 (4-1BB), which is associated with higher exhaustion marker expression and poorer patient survival in clear cell renal cell carcinoma^116,117^.

Two Tex subsets were identified from our integration of γδ T cells. Both subsets exhibited a range of exhaustion markers, including PDCD1 (PD-1), CTLA4, HAVCR2 (TIM-3), LAG3, and TIGIT (Figure 2B). Despite these markers, cytotoxic molecules such as GZMB, NKG7, PRF1, and IFNG persisted in these cells. Tex 1 cells are actively proliferating, indicated by their expression of MKI67, TOP2A, and other cell cycle-related genes such as TUBA1B, MCM3, MCM5, and MCM6. In contrast, Tex 2 cells uniquely expressed IKZF2 (Helios), a transcription factor found in Tregs, where its deficiency downregulates Foxp3 and adopts a more effector-like function^118^, specific to murine intraepithelial γδ T cells^119^. It also shows increased levels of ZNF683 (Hobit) and PRDM1 (Blimp-1), transcription factors associated with effector function and exhaustion. Hobit, the homolog of Blimp-1, is shown in humans to necessarily and sufficiently induce IFN-γ expression but has no apparent effect on Granzyme B expression^120^. Together, Hobit and Blimp-1 can regulate the TRM fate selection of Teff cells by retaining the TRM precursors in the tissue at the early stage of infection^121^.

Although Tex 2 cells lack FOXP3 and IL2RA transcription, they resemble Tregs in their high expression of CTLA4, a well-studied inhibitory receptor in the CD28 immunoglobulin subfamily. Despite retaining GZMB and IFNG transcription, which suggests potential immunosuppressive roles, Tex 2 cells also show strong expression of HLA-DR molecules such as HLA-DRA, HLA-DRB5, and HLA-DRQ. This aligns with the description of CD8^+^ HLA-DR^+^ regulatory T cells observed in human PBMCs^122^, which exhibit increased frequencies of IFN-γ and TNFα positive cells and higher degranulation after stimulation, contradicting the notion that they are functionally impaired exhausted T cells. However, CD8^+^ HLA-DR^+^ regulatory T cells reported by Machicote et al. are TIM-3 negative, whereas Tex 2 cells strongly express all exhaustion markers^123^. Thus, we conclude that Tex 2 cells are indeed exhausted T cells with some regulatory functions.

### Non-Exhausted Tumor-Infiltrating γδ T Cells Produce More Cytotoxic Molecules Than Their Normal Area Counterparts, Indicating They Act at the Frontier of Anti-Tumor Immunity

Our data integration workflow removed variation among studies while preserving biological differences between tumor-infiltrating γδ T cells and normal area γδ T cells (Figure 2A i). Composition-wise, the CTLA4^high^ Tex 2 subset is highly specific to tumors (Figure 2A iii). Differential gene expression analysis (Figure 2D) revealed that tumor-infiltrating γδ T cells, regardless of subtype, exhibited higher expression of KLF2 and GZMB. KLF2 inhibits tumor cell growth and migration in hepatocellular carcinoma (HCC), non-small cell lung cancer (NSCLC), and clear cell renal cell carcinoma (ccRCC)^124^. In T cells, as previously discussed, KLF2 regulates circulation and tissue trafficking. The higher KLF2 expression may indicate ex-circulating properties of γδ tumor-infiltrating lymphocytes (TILs), suggesting they are more likely recruited from circulation rather than migrating from adjacent tissue as TRM.

For both subsets of γδ TRM, ZNF683 (Hobit), CD81, CD82, and IFNG expression is higher in the tumor compared to normal tissue (Figure 2D). Hobit regulates IFNG production in humans, and TILs co-expressing CD81 and CD82 are shown to have higher T-cell activation and cytokine production in the NSCLC TME. Taken together, tumor-infiltrating γδ TRM has enhanced effector functions^125^. Interestingly, IL7R^-^ γδ TRM upregulates inhibitory molecules CTLA4 and PDCD1 in the TME more than IL7R^+^ γδ TRM, indicating that IL7R^+^ TRM may sustain longer life and provide a longer-lasting anti-tumor effect. Indeed, Poon et al.’s 37-marker panel CyTOF on CCR7^-^ CD45RA^-^CD69^+^ TRM shows that IL7R^-^ TRM has higher expression of PDCD1 and TIGIT^109^.

When we compared tumor Teff 1 with normal Teff 1, we observed an increased level of GNLY, GZMB, and IFNG was accompanied by higher HAVCR2 (TIM-3) (Figure 2D). This is reasonable, as T cells exhibit increased effector functions, including IFNγ and granzyme B production, as they progress towards exhaustion. Notably, for TRM subsets, there is a higher expression of ZG16 in normal tissue compared to tumors. ZG16, the human zymogen granule protein 16, is mainly expressed by mucus-secreting cells^126,127^. As a secreted protein, ZG16 has been found to directly inhibit PD-L1 and promote NK cell survival and proliferation^128^.

### IL-17-Producing Cells in Colorectal Cancer Are Mainly CD4 Helper T Cells

Though our analysis indicated γδ T cells do not produce IL-17 in human CRC, there were T cells in our integrated CRC scRNA-seq dataset that did produce IL-17. Thus, we sought to characterize these IL-17 producing cells in CRC by re-analyzing the 15 published human CRC whole-tissue scRNA-seq datasets (Figure 3A, Figure S3B). We selected cell clusters enriched with IL-17A-F transcripts and/or RORC and integrated them (Figure S4A). We identified 22,512 cells from the neoplastic and adjacent normal tissues of 187 human CRC patients across 9 studies (Figure 3A). These IL-17 producers can be categorized into seven subsets: CD4 FOXP3^+^ regulatory T cells (Treg), CD4 central memory T cells (TCM) that expressed CCR7, SELL, IL7R, S1PR1, KLF2, and TCF7, CD4 and CD8 TRM that did not express the aforementioned circulating markers but are IL7R^+^, ITGAE^+^, and ITGA1^+^. The remaining subsets were MAIT cells that had invariant TRAV1-2 usage, and ILC3 cells that expressed little CD3 and uniquely expressed KIT (Figure 3B).

**Figure 3:**
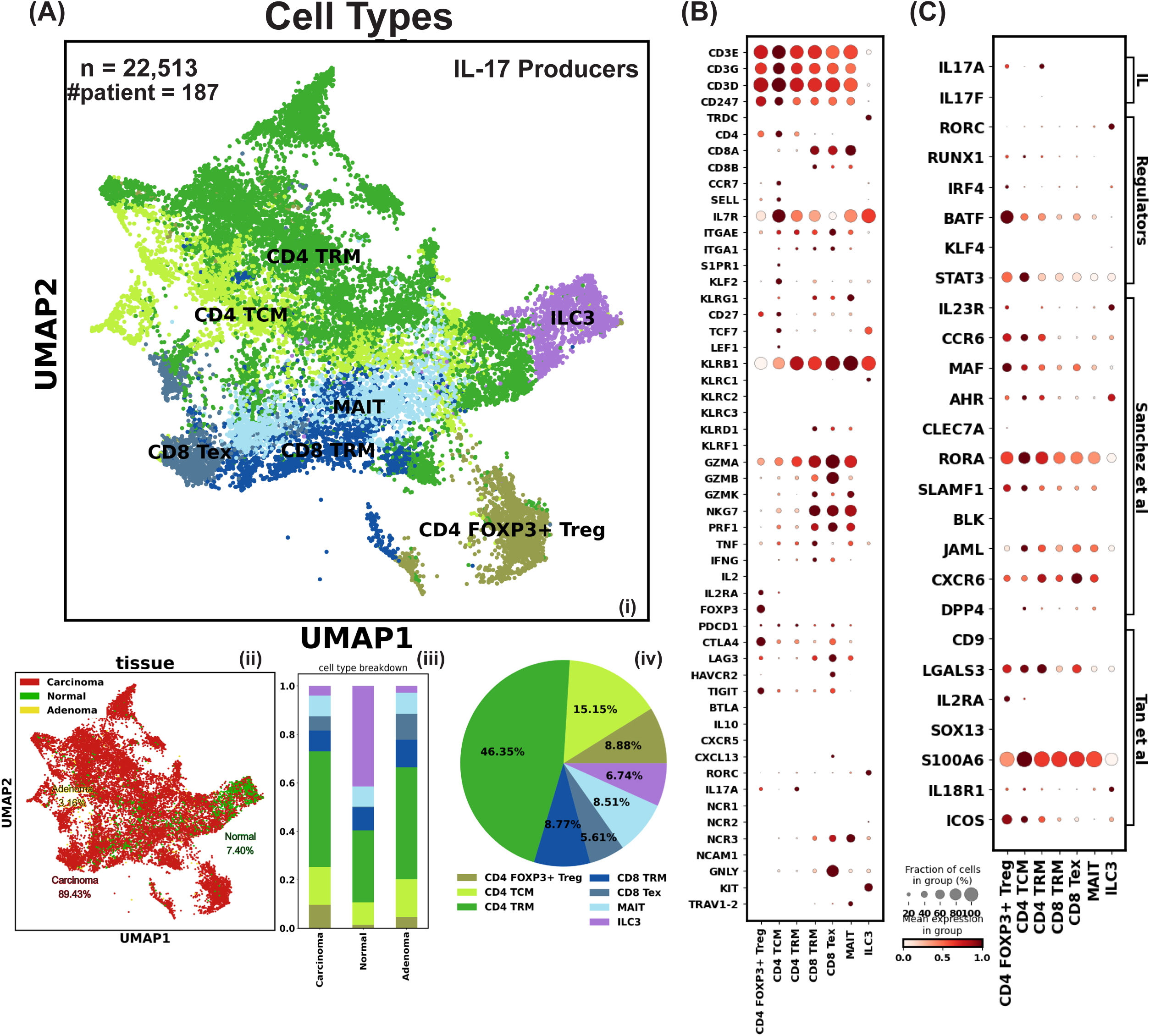
Characterization of IL-17 Producers in Human Colorectal Cancer. A) i) Uniform manifold approximation and projection (UMAP) of IL-17 producer cells showing subsets: Treg (regulatory T cells), TCM (central memory T cells), TRM (tissue-resident memory T cells), MAIT (mucosal-associated invariant T cells), Tex (exhausted T cells), and ILC3 (type 3 innate lymphoid cells). ii) UMAP of IL-17 producer cells colored by tissue origin. iii) Composition of IL-17 producer cell subtypes identified from different tissue types. iv) Composition of IL-17 producer cell subtypes in the integrated dataset. B) Dot plot displaying the level and percentage of each IL-17 producer cell subtype expressing key T cell features related to identity, mobility, tissue residency, cytotoxicity, and exhaustion. C) Dot plot showing the level and percentage of each IL-17 producer cell subtype expressing IL-17 production-related features. IL: interleukin, direct IL-17 transcript; Regulators: proteins regulating IL-17 transcription; Sanchez et al.: IL-17 producing features as defined by Sanchez Sanchez et al.; Tan et al.: IL-17 producing features from Tan et al.’s mouse study, with CD9 and LGALS3 signatures used by Reis et al. to annotate their data.

We noticed that ILC3 also has TRDC expression, and to investigate whether those cells are special γδ T or intrinsically have the TRDC expression just like NK cells, we examined an ILC scRNA-seq dataset^86^ and confirmed the TRDC expression (Figure S2C). All seven subsets broadly and highly express most of the three categories of IL-17 production features summarized earlier, which strongly contrasts with γδ T cells, further implying that human CRC-associated γδ T cells do not have high IL-17 production potential (Figure 3C). Most of these IL-17-producing cells were found in tumor tissue rather than normal tissue in CRC patients, suggesting that IL-17 production is not maintained at baseline levels and is specific to inflammation in the human colon. Based on these data, we assessed that in carcinoma tissue, more than 70% of the potential IL-17 producers were CD4 helper T cells (Figure 3A iii).

## Discussion

Our study assessed the largest number of γδ T cells to data by pooling data from nine different human CRC scRNA-seq studies. We showed that γδ T cells in human CRC tumors exhibited high cytotoxicity-related transcription both in tumors and adjacent normal areas. These cells could be further classified into Teff, TRM, Tpex, and Tex subsets. Critically, none of these subsets showed signs of active IL-17 production or production potential. Our results indicate that in the controversy over the pro-tumor effects of γδ T cells, IL-17 secreting γδ T cells are likely absent in human tumors and would not exert pro-tumor effects in human CRC as has been previously reported. Indeed, our findings strongly show that γδ T cells in human CRC are a heterogeneous population with many nuanced functions and subpopulations that overall exert anti-tumor cytotoxic effects.

Although we have presented evidence supporting the use of TRDC as a specific marker for γδ T cell identification in scRNA-seq, it is possible that Reis et al.’s TCRγδ^+^TRDC^-^TRGC1^-^TRGC2^-^ cells with marginal IL-17A and RORC transcripts but no productive chain captured in the scTCR-seq (i.e., cluster 4) represented an uncommon subset of γδ T cells with downregulated TCR transcription. Given that the current TCR-seq technique involves RNA sequencing that primes RNA containing TRGC/TRDC regions, it is possible that cells with little TRDC transcription or captured productive γδ TCR chain are transcriptionally inactive γδ T cells. This hypothesis needs further investigation, but if true, identifying such γδ17 in scRNA-seq without pre-sequencing TCRγδ gating would be very difficult given its resemblance to TCRαβ CD4 helper T cells without any γδ TCR gene module expression.

It is not clear why Wu et al. found such abundant γδ T17 cells in their human samples if not due to biological differences among patients from different cohorts. They did not show their TCRγδ gating strategy in their 2014 publication, raising questions about whether the IL-17 producers they identified are indeed γδ T cells. Rulan Ma et al. attributed the discrepancy between Wu et al. and Meraviglia et al. to “some unknown inhibitory components in the local TME” that suppress IL-17 production^50^. However, our integrative analysis of γδ T cells from multiple sites of the colon in different patients from different cohorts showed no sign of IL-17 production, which offers a refutation of the hypothesis that the absence of IL-17 is due to TME-specific conditions and reinforces the conclusion that most CRC γδ T cells do not generally exhibit such behavior.

Beyond the debate over whether human γδ T cells have a high potential for IL-17 production like their murine counterparts, McKenzie et al. suggested that human γδ17 cells may not be defined by IL-17 production but by “homing molecules, activation markers, and other subset-specific surface markers” that shape their function^129^. Nevertheless, IL-17 is an important cytokine involved in various immune processes. If its production indeed differs between human and mouse γδ T cells, such cross-species discrepancies should be carefully addressed when performing experiments on mouse models to gain insights into human γδ T cells.

Indeed, mice models are essential for studying γδ T cells and testing γδ T cell-based tumor immunotherapy, but their immunology, particularly γδ T cell development and function, differs significantly from humans^130^. In humans, CD3γ deficiency still allows γδ T cell development due to substitution by CD3δ, whereas mice lacking CD3γ exhibit a block in γδ T cell development as their γδ TCR does not incorporate CD3δ^131^. This difference results in distinct functional categorizations of mouse γδ T cells, such as IFN-γ-producing and IL-17-producing cells, each with specific expression markers and tissue localizations. In contrast, human γδ T cells display broader functional plasticity and are involved in more diverse roles^132^. Therefore, caution is warranted in interpreting results from mice models, as mouse γδ T cell responses, especially cytokine production, may not accurately reflect human γδ T cell responses, despite experimental and computational methods developed to address these discrepancies^133–137^.

## Method

### Data Collection

Human CRC whole-tissue scRNA-seq datasets were obtained from Gene Expression Omnibus (GEO) Series GSE178341^138^, GSE200997^139^, GSE221575^140^, GSE232525^141^, GSE245552^142^, GSE231559^143^, GSE188711^144^, GSE161277^145^, GSE183916^146^, GSE201348^147^, GSE108989^148^, GSE146771^149^, GSE178318^150^, and PubMed PMID35773407^151^. Reis et al.’s data and metadata were obtained from GSE205720^60^. scRNA-seq with paired scTCR-seq data on anti-CD3 gated cells used as a reference in our integrative study was obtained from Gheradin et al.^17^, GSE223809^93^. scRNA-seq of ILC3 was obtained from GSE150050^86^. Metadata that documents cells’ donors’ gender, age, CRC stage, and tissue origin, if not provided explicitly, was obtained from the supplementary information of the original paper.

### Quality Control and Pre-Processing

Only cells in the colon mucosa were kept. If processed data is not provided, Seurat^152^ and Scanpy^153^ were used for downstream analysis. For quality control, low-quality cells were dropped based on their low UMI counts (<500), high mitochondrial gene counts (>20%), and a low number of uniquely expressed genes (<200). If the unprocessed data has multiple samples, data integration was done by using Seurat. The top 3000 genes that were the most variable in as many samples as possible were used as anchors to integrate them. 3000 variable features were calculated for the whole dataset and used to perform the principal component analysis (PCA, 50 pcs). Leiden clustering was performed based on the computed neighborhood graph of observations (UMAP, 50 pcs, size of neighborhood equals 15 cells) to reveal the general subtypes.

### γδ T cells and IL-17 Producing Cells Selection and Integration

In each dataset, cell clusters that highly co-express CD3D, CD3E, CD3G, CD247, and TRDC were identified as γδ T and cell clusters that express either IL17A-F or RORC were identified as potential IL-17 producers. γδ clusters with more than 50 cells were identified from GSE161277, GSE245552, GSE200997, GSE178341, GSE221575, GSE178318, GSE231559, GSE188711, and PubMed PMID35773407. IL-17 producer clusters with more than 50 cells were identified from GSE245552, GSE178318, GSE231559, GSE161277, GSE178341, GSE108989, GSE188711, GSE200997, and PubMed PMID35773407. Raw counts of cells in the γδ clusters of all studies were normalized by SCTransform^154^ and integrated using Seurat using 3000 variable features as anchors.

### Differentially Expressed Gene Analysis

Differentially expressed gene (DEG) analysis was performed using Monocle 3^155^. This involved fitting a generalized linear model (GLM) to SCTransform-corrected counts, employing a quasi-Poisson distribution to account for the mean and variance in the data. The model assessed gene expression levels of TRM and Teff, respectively, across tissue origins. The calculation procedure is described in our previous work^156^. Briefly, for a given gene, let *y_ij_* represent the observed expression level for cell *i* in condition *j*, for cell *i* in condition *j*, where *i* = 1, 2,…, n, and *j* = 1, 2 (two origins, tumor or adjacent normal tissue). The quasi-Poisson GLM can be written as *Y_ij_* ∼ *QuasiPoisson*(*μ_ij_*, *ϕ*), *log*(*μ_ij_*) = *X_ij_β_j_*, where *μ_ij_* is the expected expression level for cell *i* in condition j, *ϕ* is the dispersion parameter, **X*_ij_* is the design matrix representing covariates (e.g., experimental conditions, batch effects), and *β_j_* is the vector of regression coefficients for condition *j*. For each gene, the GLM is fit to the data using maximum likelihood estimation, which involves finding the *β_j_* and *ϕ* values that maximize the likelihood of the observed data. A likelihood ratio test is performed to test for differential expression between the two conditions. This compares the likelihood of the data under the full model (with separate *β_j_* values for each condition) to the likelihood under the null model (with the same *β_j_* value for both conditions). Genes with q-values below 0.05 and a log2 fold change larger than 1 were considered differentially expressed.

## Code Availability

The code used for processing datasets can be found at https://github.com/Brubaker-Lab/CRCgdT.

## Data Availability

All data necessary to reproduce the findings of this manuscript are available within their originally published studies as cited in the text.

## Author contributions statement

Conceptualization: R.R. D.B.; Methodology: R.R., D.B.; Analysis: R.R.; Writing: R.R., M.T. D.B.; Administration: D.B.

---

**Figure S1:**
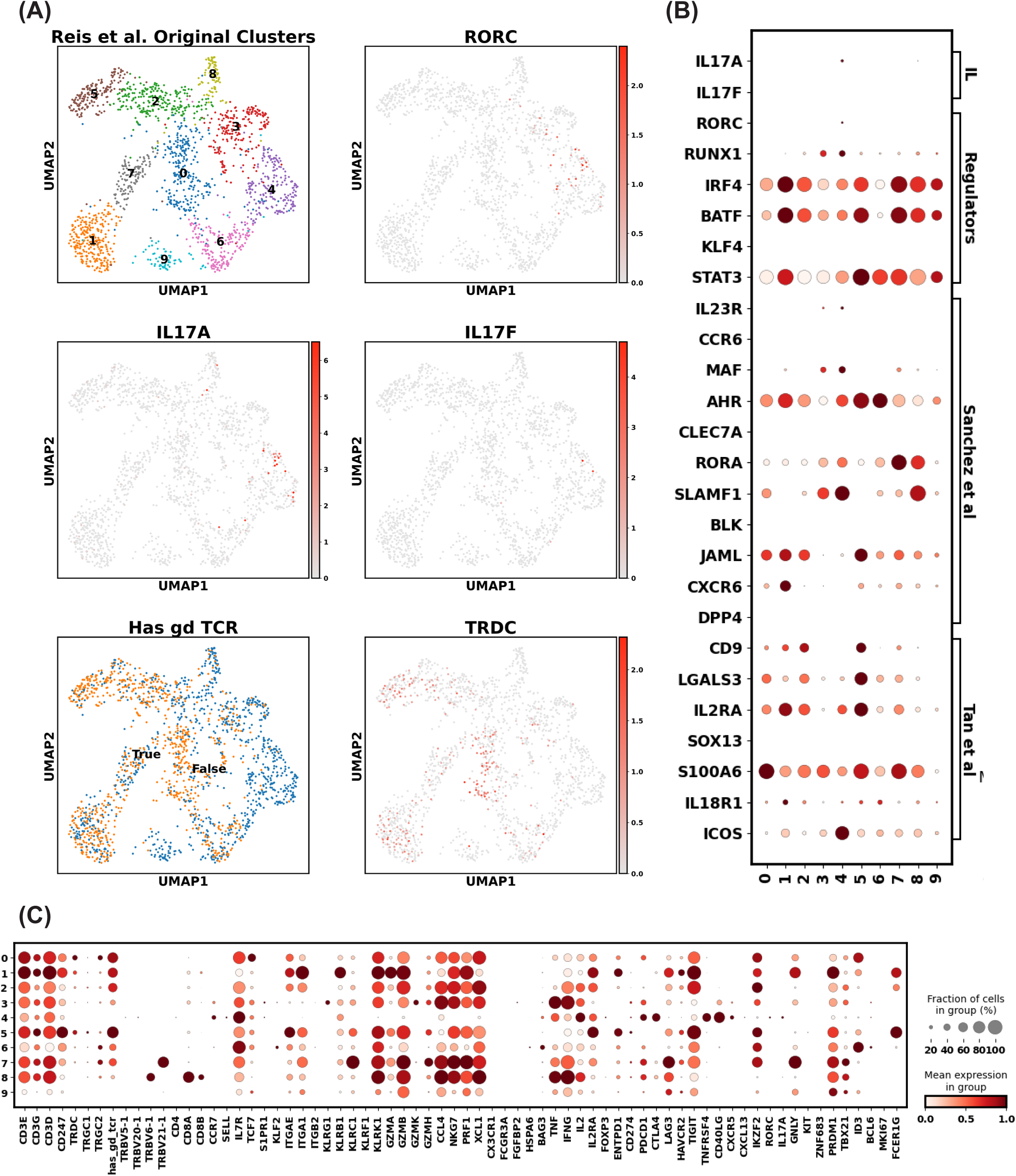
Query Gene Expression in Reis et al.’s Human Colorectal Cancer γδ T Cells Data. A) Reproduction of the uniform manifold approximation and projection (UMAP) of Reis et al.’s γδ T cells showing cell clusters, expression levels of RORC, IL17A, IL17F, TRDC, and binary labeling of TCRγδ sequencing status. Coordinates and cluster information were provided by Reis et al. B) Dot plot displaying the level and percentage of each cell cluster expressing key T cell features related to identity, mobility, tissue residency, cytotoxicity, and exhaustion. C) Dot plot showing the level and percentage of each cell cluster expressing IL-17 production-related features. IL: interleukin, direct IL-17 transcript; Regulators: proteins regulating IL-17 transcription; Sanchez et al.: IL-17 producing features as defined by Sanchez Sanchez et al.; Tan et al.: IL-17 producing features from Tan et al.’s mouse study, with CD9 and LGALS3 signatures used by Reis et al. to annotate their data.

**Figure S2:**
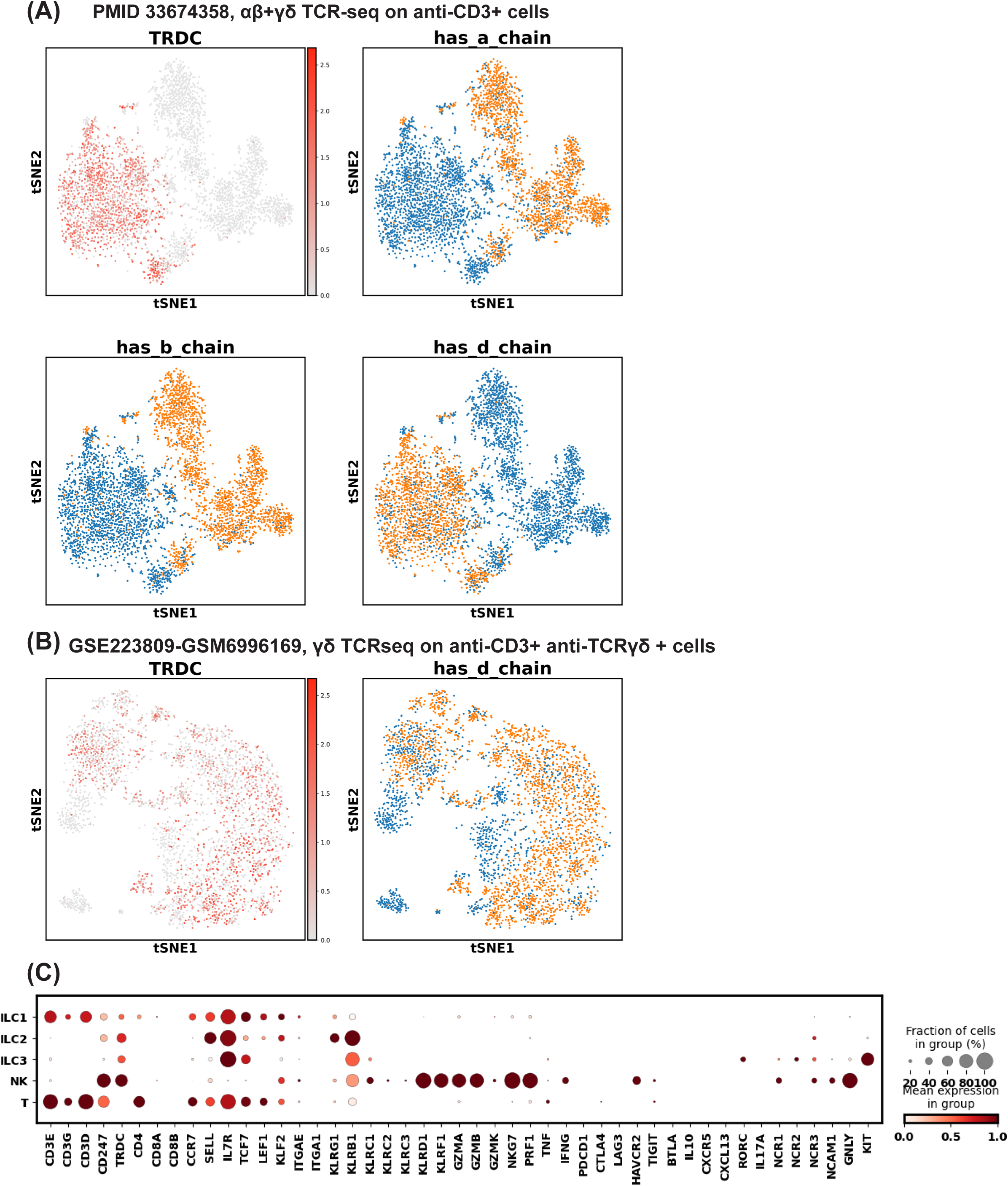
Datasets Used as Reference to Establish Reliable γδ T Cells Transcriptomic Markers. A) Uniform manifold approximation and projection (UMAP) of T cells in the study PMID33674358 colored by TRDC expression and whether a TRA/TRB/TRD chain is sequenced in the cell. B) UMAP of flow-sorted CD3^+^TCRγδ^+^ T cells in the study GSE223809 colored by TRDC expression and whether a TRD chain is sequenced in the cell. C) Dot plot showing the level and percentage of each cell type expressing key features that delineate T, NK, and ILC in the study GSE150050. Cell type annotation is provided by the original authors.

**Figure S3:**
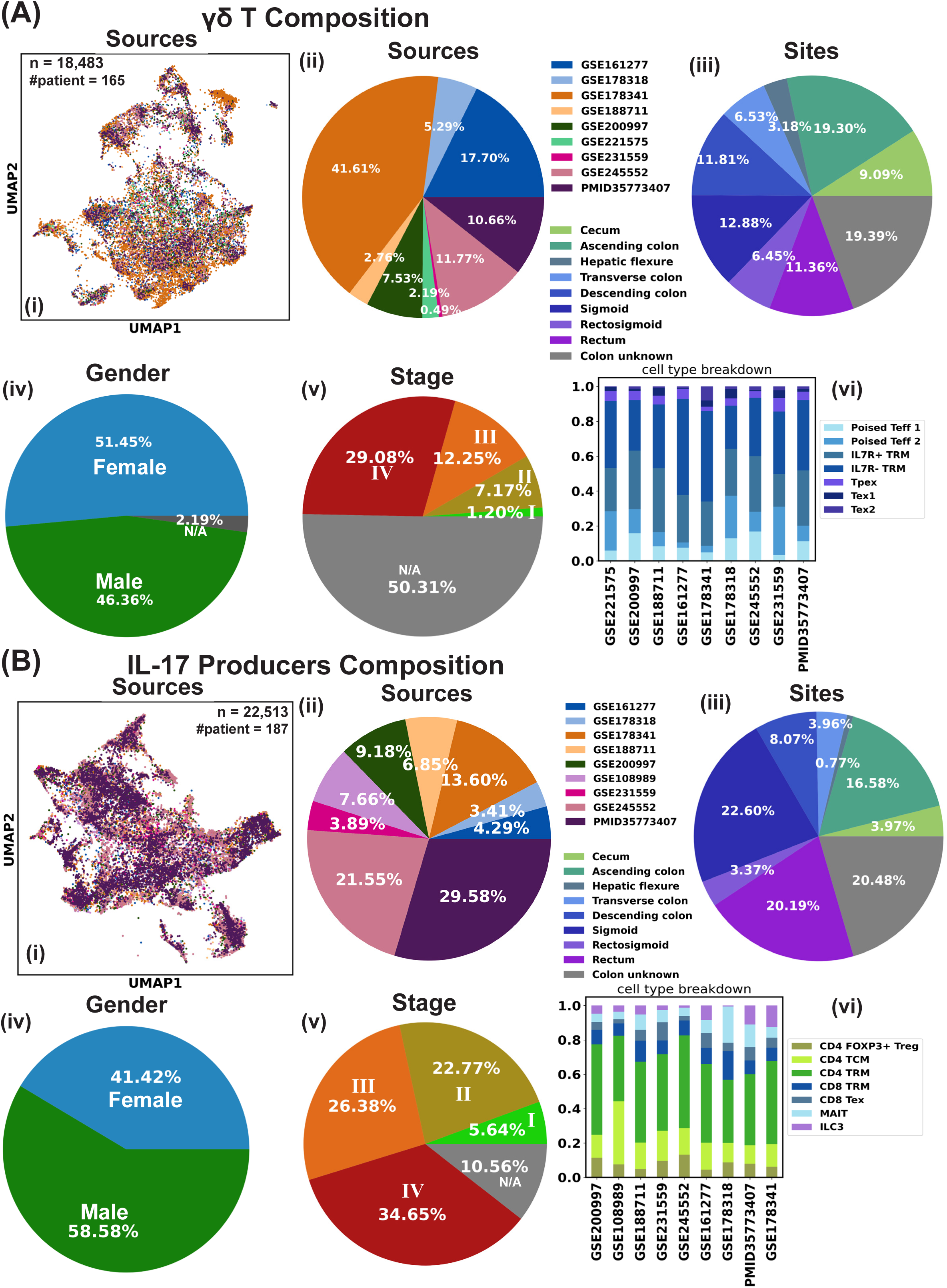
Data Composition Overview. A) i) Uniform manifold approximation and projection (UMAP) of γδ T cells colored by data sources. ii) Pie chart showing the data source contribution to the integrated data. iii) Pie chart showing the composition of γδ T cells’ site origins. iv) Pie chart showing the composition of γδ T cells’ donors’ gender. v) Pie chart showing the composition of γδ T cells’ donors’ CRC stage. For cells from donors with unknown stage or cells from the adjacent normal tissue or adenoma, N/A is assigned. vi) Bar plot showing the data source contribution to each identified γδ T cell subtype. B) i) Uniform manifold approximation and projection (UMAP) of IL-17 Producers colored by data sources. ii) Pie chart showing the data source contribution to the integrated datas. iii) Pie chart showing the composition of IL-17 Producers’ site origins. iv) Pie chart showing the composition of IL-17 Producers’ donors’ gender. v) Pie chart showing the composition of IL-17 Producers’ donors’ CRC stage. For cells from donors with unknown stage or cells from the adjacent normal tissue or adenoma, N/A is assigned. vi) Bar plot showing the data source contribution to each identified γδ T cell subtype.

**Figure S4:**
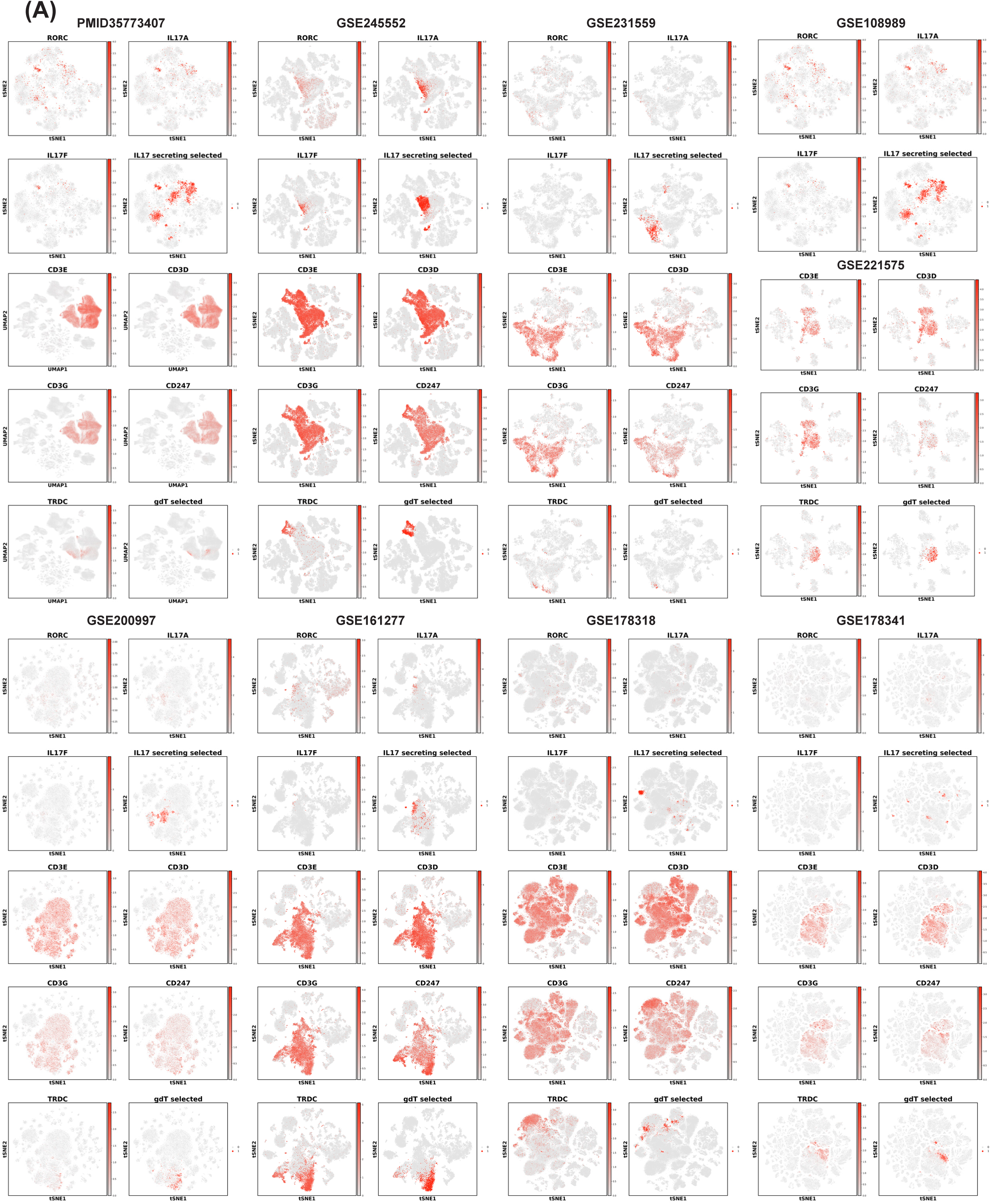
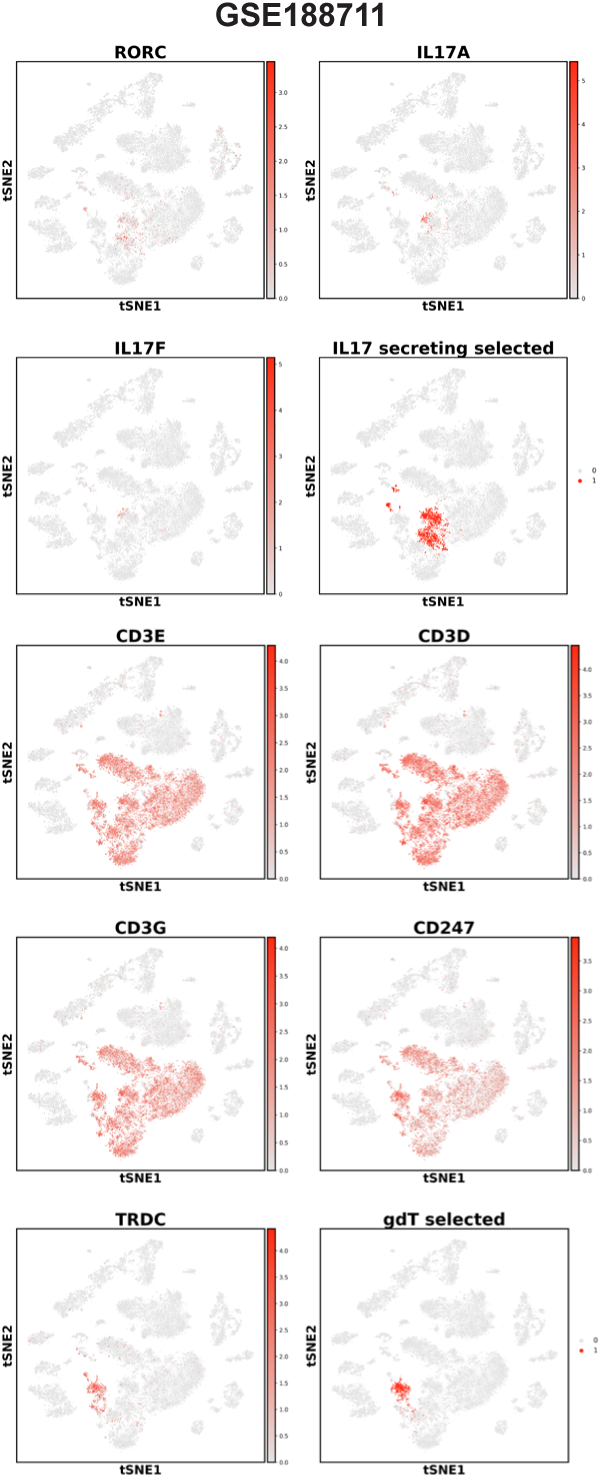
Details of the γδ T Cells and IL-17 Producer Cells Selection. A) Uniform manifold approximation and projection (UMAP) of T cells in all individual studies used, colored by expression of CD3D, CD3E, CD3G, TRDC, RORC, IL17A, and IL17F, along with whether a cell has been selected as a γδ T cell or an IL-17 producer.

## Reference

1. Attaf, M., Legut, M., Cole, D. K. & Sewell, A. K. The T cell antigen receptor: the Swiss army knife of the immune system. Clin. Exp. Immunol. 181, 1–18 (2015).

2. Bonneville, M., O’Brien, R. L. & Born, W. K. Gammadelta T cell effector functions: a blend of innate programming and acquired plasticity. Nat. Rev. Immunol. 10, 467–478 (2010).

3. Chodaczek, G., Papanna, V., Zal, M. A. & Zal, T. Body-barrier surveillance by epidermal γδ TCRs. Nat. Immunol. 13, 272–282 (2012).

4. Edelblum, K. L. et al. Dynamic migration of γδ intraepithelial lymphocytes requires occludin. Proc. Natl. Acad. Sci. 109, 7097–7102 (2012).

5. Cheng, M. & Hu, S. Lung-resident γδ T cells and their roles in lung diseases. Immunology 151, 375–384 (2017).

6. Matsukawa, M., Kumamoto, Y., Hirose, T. & Matsuura, A. [Tissue gamma/delta T cells in experimental urinary tract infection relationship between other immuno-competent cells]. Kansenshogaku Zasshi 68, 1498–1511 (1994).

7. Salim, M. et al. BTN3A1 Discriminates γδ T Cell Phosphoantigens from Nonantigenic Small Molecules via a Conformational Sensor in Its B30.2 Domain. ACS Chem. Biol. 12, 2631– 2643 (2017).

8. Uldrich, A. P. et al. CD1d-lipid antigen recognition by the γδ TCR. Nat. Immunol. 14, 1137–1145 (2013).

9. Silva-Santos, B. & Strid, J. Working in “NK Mode”: Natural Killer Group 2 Member D and Natural Cytotoxicity Receptors in Stress-Surveillance by γδ T Cells. Front. Immunol. 9, (2018).

10. Dong, R., Zhang, Y., Xiao, H. & Zeng, X. Engineering γδ T Cells: Recognizing and Activating on Their Own Way. Front. Immunol. 13, 889051 (2022).

11. Harly, C., et al. Human γδ T cell sensing of AMPK-dependent metabolic tumor reprogramming through TCR recognition of EphA2. Sci. Immunol. 6, eaba9010 (2021).

12. Vantourout, P. & Hayday, A. Six-of-the-best: unique contributions of γδ T cells to immunology. Nat. Rev. Immunol. 13, 88–100 (2013).

13. Ribot, J. C., Lopes, N. & Silva-Santos, B. γδ T cells in tissue physiology and surveillance. Nat. Rev. Immunol. 21, 221–232 (2021).

14. Rong, L. et al. Analysis of tumor-infiltrating gamma delta T cells in rectal cancer. World J. Gastroenterol. 22, 3573–3580 (2016).

15. Patin, E. C. et al. Type I IFN Receptor Signaling Controls IL7-Dependent Accumulation and Activity of Protumoral IL17A-Producing γδT Cells in Breast Cancer. Cancer Res. 78, 195– 204 (2018).

16. Nezhad Shamohammadi, F., et al. Controversial role of γδ T cells in pancreatic cancer. Int. Immunopharmacol. 108, 108895 (2022).

17. Gherardin, N. A. et al. γδ T Cells in Merkel Cell Carcinomas Have a Proinflammatory Profile Prognostic of Patient Survival. Cancer Immunol. Res. 9, 612–623 (2021).

18. Meraviglia, S. et al. Distinctive features of tumor-infiltrating γδ T lymphocytes in human colorectal cancer. OncoImmunology (2017).

19. Amicarella, F. et al. Dual role of tumour-infiltrating T helper 17 cells in human colorectal cancer. Gut 66, 692–704 (2017).

20. Altvater, B. et al. Activated human γδ T cells induce peptide-specific CD8+ T-cell responses to tumor-associated self-antigens. Cancer Immunol. Immunother. CII 61, 385–396 (2011).

21. Kunzmann, V., Bauer, E., Feurle, J., Tony, F. W., Hans-Peter & Wilhelm, M. Stimulation of γδ T cells by aminobisphosphonates and induction of antiplasma cell activity in multiple myeloma. Blood 96, 384–392 (2000).

22. Viey, E. et al. Phosphostim-activated gamma delta T cells kill autologous metastatic renal cell carcinoma. J. Immunol. Baltim. Md 1950 174, 1338–1347 (2005).

23. Mattarollo, S. R., Kenna, T., Nieda, M. & Nicol, A. J. Chemotherapy and zoledronate sensitize solid tumour cells to Vgamma9Vdelta2 T cell cytotoxicity. Cancer Immunol. Immunother. CII 56, 1285–1297 (2007).

24. Tokuyama, H. et al. V gamma 9 V delta 2 T cell cytotoxicity against tumor cells is enhanced by monoclonal antibody drugs--rituximab and trastuzumab. Int. J. Cancer 122, 2526– 2534 (2008).

25. Todaro, M. et al. Efficient killing of human colon cancer stem cells by gammadelta T lymphocytes. J. Immunol. Baltim. Md 1950 182, 7287–7296 (2009).

26. D’Asaro, M. et al. V gamma 9V delta 2 T lymphocytes efficiently recognize and kill zoledronate-sensitized, imatinib-sensitive, and imatinib-resistant chronic myelogenous leukemia cells. J. Immunol. Baltim. Md 1950 184, 3260–3268 (2010).

27. Maniar, A. et al. Human γδ T lymphocytes induce robust NK cell–mediated antitumor cytotoxicity through CD137 engagement. Blood 116, 1726–1733 (2010).

28. Fisher, J. P. H. et al. Neuroblastoma killing properties of Vδ2 and Vδ2-negative γδT cells following expansion by artificial antigen-presenting cells. Clin. Cancer Res. Off. J. Am. Assoc. Cancer Res. 20, 5720–5732 (2014).

29. Holmen Olofsson, G., et al. Vγ9Vδ2 T Cells Concurrently Kill Cancer Cells and Cross-Present Tumor Antigens. Front. Immunol. 12, 645131 (2021).

30. Wen, L. et al. Germinal center formation, immunoglobulin class switching, and autoantibody production driven by ‘non alpha/beta’ T cells. J. Exp. Med. 183, 2271–2282 (1996).

31. Girardi, M. et al. Characterizing the Protective Component of the αβ T Cell Response to Transplantable Squamous Cell Carcinoma. J. Invest. Dermatol. 122, 699–706 (2004).

32. Gao, Y. et al. γδ T Cells Provide an Early Source of Interferon γ in Tumor Immunity. J. Exp. Med. 198, 433–442 (2003).

33. Street, S. E. A. et al. Innate Immune Surveillance of Spontaneous B Cell Lymphomas by Natural Killer Cells and γδ T Cells. J. Exp. Med. 199, 879–884 (2004).

34. Riond, J., Rodriguez, S., Nicolau, M.-L., al Saati, T. & Gairin, J. E. In vivo major histocompatibility complex class I (MHCI) expression on MHCIlow tumor cells is regulated by γδ T and NK cells during the early steps of tumor growth. Cancer Immun. J. Acad. Cancer Immunol. 9, 10 (2009).

35. Capietto, A.-H., Martinet, L. & Fournié, J.-J. Stimulated γδ T cells increase the in vivo efficacy of trastuzumab in HER-2+ breast cancer. J. Immunol. Baltim. Md 1950 187, 1031–1038 (2011).

36. He, W. et al. Naturally Activated Vγ4 γδ T Cells Play a Protective Role in Tumor Immunity Through Expression of Eomesodermin. J. Immunol. Baltim. Md 1950 185, 10.4049/jimmunol.0903767 (2010).

37. Lança, T. Resposta dos linfócitos T ɣδ a tumoures : recrutamento, reconhecimento e funções. (2013).

38. Dalessandri, T., Crawford, G., Hayes, M., Castro Seoane, R. & Strid, J. IL-13 from intraepithelial lymphocytes regulates tissue homeostasis and protects against carcinogenesis in the skin. Nat. Commun. 7, 12080 (2016).

39. Rezende, R. M. et al. γδ T cells control humoral immune response by inducing T follicular helper cell differentiation. Nat. Commun. 9, 3151 (2018).

40. Caccamo, N. et al. CXCR5 identifies a subset of Vgamma9Vdelta2 T cells which secrete IL-4 and IL-10 and help B cells for antibody production. J. Immunol. Baltim. Md 1950 177, 5290–5295 (2006).

41. de Vries, N. L. et al. γδ T cells are effectors of immunotherapy in cancers with HLA class I defects. Nature 613, 743–750 (2023).

42. Pansy, K. et al. Immune Regulatory Processes of the Tumor Microenvironment under Malignant Conditions. Int. J. Mol. Sci. 22, 13311 (2021).

43. Tawfik, D. et al. TRAIL-Receptor 4 Modulates γδ T Cell-Cytotoxicity Toward Cancer Cells. Front. Immunol. 10, 2044 (2019).

44. Kärre, K., Ljunggren, H. G., Piontek, G. & Kiessling, R. Selective rejection of H-2-deficient lymphoma variants suggests alternative immune defence strategy. Nature 319, 675–678 (1986).

45. World Health Organization. Cancer. World Health Organization https://www.who.int/news-room/fact-sheets/detail/cancer.

46. Keum, N. & Giovannucci, E. Global burden of colorectal cancer: emerging trends, risk factors and prevention strategies. Nat. Rev. Gastroenterol. Hepatol. 16, 713–732 (2019).

47. Curry, W. J. et al. Academic detailing to increase colorectal cancer screening by primary care practices in Appalachian Pennsylvania. BMC Health Serv. Res. 11, 112 (2011).

48. Hossain, Md. S., et al. Colorectal Cancer: A Review of Carcinogenesis, Global Epidemiology, Current Challenges, Risk Factors, Preventive and Treatment Strategies. Cancers 14, 1732 (2022).

49. Ryuk, J. P. et al. Predictive factors and the prognosis of recurrence of colorectal cancer within 2 years after curative resection. Ann. Surg. Treat. Res. 86, 143–151 (2014).

50. Ma, R., Yuan, D., Guo, Y., Yan, R. & Li, K. Immune Effects of γδ T Cells in Colorectal Cancer: A Review. Front. Immunol. 11, (2020).

51. Guzman, G., Reed, M. R., Bielamowicz, K., Koss, B. & Rodriguez, A. CAR-T Therapies in Solid Tumors: Opportunities and Challenges. Curr. Oncol. Rep. 25, 479–489 (2023).

52. Maalej, K. M. et al. CAR-cell therapy in the era of solid tumor treatment: current challenges and emerging therapeutic advances. Mol. Cancer 22, 20 (2023).

53. Rischer, M. et al. Human gammadelta T cells as mediators of chimaeric-receptor redirected anti-tumour immunity. Br. J. Haematol. 126, 583–592 (2004).

54. Deniger, D. C. et al. Bispecific T-cells expressing polyclonal repertoire of endogenous γδ T-cell receptors and introduced CD19-specific chimeric antigen receptor. Mol. Ther. J. Am. Soc. Gene Ther. 21, 638–647 (2013).

55. Rozenbaum, M. et al. Gamma-Delta CAR-T Cells Show CAR-Directed and Independent Activity Against Leukemia. Front. Immunol. 11, 1347 (2020).

56. Zhai, X. et al. MUC1-Tn-targeting chimeric antigen receptor-modified Vγ9Vδ2 T cells with enhanced antigen-specific anti-tumor activity. Am. J. Cancer Res. 11, 79–91 (2021).

57. Nagai, N., Kudo, Y., Aki, D., Nakagawa, H. & Taniguchi, K. Immunomodulation by Inflammation during Liver and Gastrointestinal Tumorigenesis and Aging. Int. J. Mol. Sci. 22, 2238 (2021).

58. Wu, P. et al. γδT17 cells promote the accumulation and expansion of myeloid-derived suppressor cells in human colorectal cancer. Immunity 40, 785–800 (2014).

59. Hu, G. et al. Tumor-infiltrating CD39+ γδTregs are novel immunosuppressive T cells in human colorectal cancer. OncoImmunology 6, e1277305 (2017).

60. Reis, B. S. et al. TCR-Vγδ usage distinguishes protumor from antitumor intestinal γδ T cell subsets. Science 377, 276–284 (2022).

61. Tan, L. et al. Single-Cell Transcriptomics Identifies the Adaptation of Scart1+ Vγ6+ T Cells to Skin Residency as Activated Effector Cells. Cell Rep. 27, 3657–3671.e4 (2019).

62. Sutton, C. E. et al. Interleukin-1 and IL-23 induce innate IL-17 production from gammadelta T cells, amplifying Th17 responses and autoimmunity. Immunity 31, 331–341 (2009).

63. Roark, C. L. et al. Exacerbation of Collagen-Induced Arthritis by Oligoclonal, IL-17-Producing γδ T Cells1. J. Immunol. 179, 5576–5583 (2007).

64. Sato, K. et al. Production of IL-17A at Innate Immune Phase Leads to Decreased Th1 Immune Response and Attenuated Host Defense against Infection with Cryptococcus deneoformans. J. Immunol. 205, 686–698 (2020).

65. Liu, H. et al. Staphylococcus aureus Epicutaneous Exposure Drives Skin Inflammation via IL-36-Mediated T Cell Responses. Cell Host Microbe 22, 653–666.e5 (2017).

66. Sutherland, T. E. et al. Chitinase-like proteins promote IL-17-mediated neutrophilia in a tradeoff between nematode killing and host damage. Nat. Immunol. 15, 1116–1125 (2014).

67. Sheel, M. et al. IL-17A–Producing γδ T Cells Suppress Early Control of Parasite Growth by Monocytes in the Liver. J. Immunol. 195, 5707–5717 (2015).

68. McGinley, A. M. et al. Interleukin-17A Serves a Priming Role in Autoimmunity by Recruiting IL-1β-Producing Myeloid Cells that Promote Pathogenic T Cells. Immunity 52, 342–356.e6 (2020).

69. Gil-Pulido, J. et al. Interleukin-23 receptor expressing γδ T cells locally promote early atherosclerotic lesion formation and plaque necrosis in mice. Cardiovasc. Res. 118, 2932–2945 (2022).

70. Silva-Santos, B., Mensurado, S. & Coffelt, S. B. γδ T cells: pleiotropic immune effectors with therapeutic potential in cancer. Nat. Rev. Cancer 19, 392–404 (2019).

71. Suzuki, T., Hayman, L., Kilbey, A., Edwards, J. & Coffelt, S. B. Gut γδ T cells as guardians, disruptors, and instigators of cancer. Immunol. Rev. 298, 198–217 (2020).

72. Sagar. Unraveling the secrets of γδ T cells with single-cell biology. J. Leukoc. Biol. 115, 47–56 (2024).

73. Michel, M.-L. et al. Interleukin 7 (IL-7) selectively promotes mouse and human IL-17-producing γδ cells. Proc. Natl. Acad. Sci. U. S. A. 109, 17549–17554 (2012).

74. Castillo-González, R., Cibrian, D. & Sánchez-Madrid, F. Dissecting the complexity of γδ T-cell subsets in skin homeostasis, inflammation, and malignancy. J. Allergy Clin. Immunol. 147, 2030–2042 (2021).

75. Deknuydt, F., Scotet, E. & Bonneville, M. Modulation of inflammation through IL-17 production by gammadelta T cells: mandatory in the mouse, dispensable in humans? Immunol. Lett. 127, 8–12 (2009).

76. Dupraz, L. et al. Gut microbiota-derived short-chain fatty acids regulate IL-17 production by mouse and human intestinal γδ T cells. Cell Rep. 36, (2021).

77. Fenoglio, D. et al. Vdelta1 T lymphocytes producing IFN-gamma and IL-17 are expanded in HIV-1-infected patients and respond to Candida albicans. Blood 113, 6611–6618 (2009).

78. Peng, M. et al. Interleukin 17-Producing γδ T Cells Increased in Patients with Active Pulmonary Tuberculosis. Cell. Mol. Immunol. 5, 203–208 (2008).

79. Wang, X. et al. Host-derived lipids orchestrate pulmonary γδ T cell response to provide early protection against influenza virus infection. Nat. Commun. 12, 1914 (2021).

80. Khan, D. & Ansar Ahmed, S. Regulation of IL-17 in autoimmune diseases by transcriptional factors and microRNAs. Front. Genet. 6, (2015).

81. Sanchez Sanchez, G., et al. Identification of distinct functional thymic programming of fetal and pediatric human γδ thymocytes via single-cell analysis. Nat. Commun. 13, 5842 (2022).

82. Iwakura, Y. & Ishigame, H. The IL-23/IL-17 axis in inflammation. J. Clin. Invest. 116, 1218–1222 (2006).

83. Liu, H. & Rohowsky-Kochan, C. Regulation of IL-17 in human CCR6+ effector memory T cells. J. Immunol. Baltim. Md 1950 180, 7948–7957 (2008).

84. Zuberbuehler, M. K. et al. The transcription factor c-Maf is essential for the commitment of IL-17-producing γδ T cells. Nat. Immunol. 20, 73–85 (2019).

85. McCluskey, D. et al. Single-cell analysis implicates TH17-to-TH2 cell plasticity in the pathogenesis of palmoplantar pustulosis. J. Allergy Clin. Immunol. 150, 882–893 (2022).

86. Mazzurana, L. et al. Tissue-specific transcriptional imprinting and heterogeneity in human innate lymphoid cells revealed by full-length single-cell RNA-sequencing. Cell Res. 31, 554–568 (2021).

87. Wang, X. et al. Single-Cell RNA-Seq of T Cells in B-ALL Patients Reveals an Exhausted Subset with Remarkable Heterogeneity. Adv. Sci. 8, 2101447 (2021).

88. Pizzolato, G. et al. Single-cell RNA sequencing unveils the shared and the distinct cytotoxic hallmarks of human TCRVδ1 and TCRVδ2 γδ T lymphocytes. Proc. Natl. Acad. Sci. 116, 11906–11915 (2019).

89. He, W. et al. Hepatocellular carcinoma-infiltrating γδ T cells are functionally defected and allogenic Vδ2+ γδ T cell can be a promising complement. Clin. Transl. Med. 12, e800 (2022).

90. Mullan, K. A., de Vrij, N., Valkiers, S. & Meysman, P. Current annotation strategies for T cell phenotyping of single-cell RNA-seq data. Front. Immunol. 14, (2023).

91. Boufea, K. et al. Single-cell RNA sequencing of human breast tumour-infiltrating immune cells reveals a γδ T-cell subtype associated with good clinical outcome. Life Sci. Alliance 4, (2021).

92. Song, Z. et al. Human γδ T cell identification from single-cell RNA sequencing datasets by modular TCR expression. J. Leukoc. Biol. 114, 630–638 (2023).

93. Rancan, C. et al. Exhausted intratumoral Vδ2-γδ T cells in human kidney cancer retain effector function. Nat. Immunol. 24, 612–624 (2023).

94. Boehme, L., Roels, J. & Taghon, T. Development of γδ T cells in the thymus - A human perspective. Semin. Immunol. 61–64, 101662 (2022).

95. Lai, D.-M., Shu, Q. & Fan, J. The origin and role of innate lymphoid cells in the lung. Mil. Med. Res. 3, 25 (2016).

96. Shi, M. et al. Cytoplasmic Expression of CD3ε Heterodimers by Flow Cytometry Rapidly Distinguishes Between Mature T-Cell and Natural Killer-Cell Neoplasms. Am. J. Clin. Pathol. 154, 683–691 (2020).

97. Lanier, L. L., Chang, C., Spits, H. & Phillips, J. H. Expression of cytoplasmic CD3 epsilon proteins in activated human adult natural killer (NK) cells and CD3 gamma, delta, epsilon complexes in fetal NK cells. Implications for the relationship of NK and T lymphocytes. J. Immunol. Baltim. Md 1950 149, 1876–1880 (1992).

98. Capone, A. & Volpe, E. Transcriptional Regulators of T Helper 17 Cell Differentiation in Health and Autoimmune Diseases. Front. Immunol. 11, (2020).

99. Volpe, E. et al. A critical function for transforming growth factor-beta, interleukin 23 and proinflammatory cytokines in driving and modulating human T(H)-17 responses. Nat. Immunol. 9, 650–657 (2008).

100. Ivanov, I. I. et al. The orphan nuclear receptor RORgammat directs the differentiation program of proinflammatory IL-17+ T helper cells. Cell 126, 1121–1133 (2006).

101. Manel, N., Unutmaz, D. & Littman, D. R. The differentiation of human T(H)-17 cells requires transforming growth factor-beta and induction of the nuclear receptor RORgammat. Nat. Immunol. 9, 641–649 (2008).

102. Harberts, A. et al. Interferon regulatory factor 4 controls effector functions of CD8+ memory T cells. Proc. Natl. Acad. Sci. U. S. A. 118, e2014553118 (2021).

103. Sun, Q. et al. STAT3 regulates CD8+ T cell differentiation and functions in cancer and acute infection. J. Exp. Med. 220, e20220686 (2023).

104. Kurachi, M. et al. The transcription factor BATF operates as an essential differentiation checkpoint in early effector CD8+ T cells. Nat. Immunol. 15, 373–383 (2014).

105. de Lima, K. A. et al. TGFβ1 signaling sustains aryl hydrocarbon receptor (AHR) expression and restrains the pathogenic potential of TH17 cells by an AHR-independent mechanism. Cell Death Dis. 9, 1130 (2018).

106. FitzPatrick, M. E. B. et al. Human intestinal tissue-resident memory T cells comprise transcriptionally and functionally distinct subsets. Cell Rep. 34, 108661 (2021).

107. Zhou, X. et al. HSPA6 is Correlated With the Malignant Progression and Immune Microenvironment of Gliomas. Front. Cell Dev. Biol. 10, 833938 (2022).

108. Li, H. et al. Exploring the dynamics and influencing factors of CD4 T cell activation using single-cell RNA-seq. iScience 26, 107588 (2023).

109. Poon, M. M. L. et al. Tissue adaptation and clonal segregation of human memory T cells in barrier sites. Nat. Immunol. 24, 309–319 (2023).

110. Mac Donald, A., et al. KLRC1 knockout overcomes HLA-E-mediated inhibition and improves NK cell antitumor activity against solid tumors. Front. Immunol. 14, (2023).

111. Chou, C. et al. Programme of self-reactive innate-like T cell-mediated cancer immunity. Nature 605, 139–145 (2022).

112. Utzschneider, D. T. et al. T cells maintain an exhausted phenotype after antigen withdrawal and population reexpansion. Nat. Immunol. 14, 603–610 (2013).

113. Utzschneider, D. T. et al. T Cell Factor 1-Expressing Memory-like CD8+ T Cells Sustain the Immune Response to Chronic Viral Infections. Immunity 45, 415–427 (2016).

114. Speiser, D. E. et al. T cell differentiation in chronic infection and cancer: functional adaptation or exhaustion? Nat. Rev. Immunol. 14, 768–774 (2014).

115. Im, S. J. et al. Defining CD8+ T cells that provide the proliferative burst after PD-1 therapy. Nature 537, 417–421 (2016).

116. Li, Y. et al. Tumor-infiltrating TNFRSF9+ CD8+ T cells define different subsets of clear cell renal cell carcinoma with prognosis and immunotherapeutic response. Oncoimmunology 9, 1838141.

117. van der Leun, A. M., Thommen, D. S. & Schumacher, T. N. CD8+ T cell states in human cancer: insights from single-cell analysis. Nat. Rev. Cancer 20, 218–232 (2020).

118. Lam, A. J., Uday, P., Gillies, J. K. & Levings, M. K. Helios is a marker, not a driver, of human Treg stability. Eur. J. Immunol. 52, 75–84 (2022).

119. Apostolov, A. K. et al. Common and Exclusive Features of Intestinal Intraepithelial γδ T Cells and Other γδ T Cell Subsets. ImmunoHorizons 6, 515–527 (2022).

120. Vieira Braga, F. A., et al. Blimp-1 homolog Hobit identifies effector-type lymphocytes in humans. Eur. J. Immunol. 45, 2945–2958 (2015).

121. Parga-Vidal, L. et al. Hobit and Blimp-1 regulate TRM abundance after LCMV infection by suppressing tissue exit pathways of TRM precursors. Eur. J. Immunol. 52, 1095–1111 (2022).

122. Yoshikawa, T. et al. Genetic ablation of PRDM1 in antitumor T cells enhances therapeutic efficacy of adoptive immunotherapy. Blood 139, 2156–2172 (2022).

123. Machicote, A., Belén, S., Baz, P., Billordo, L. A. & Fainboim, L. Human CD8+HLA- DR+ Regulatory T Cells, Similarly to Classical CD4+Foxp3+ Cells, Suppress Immune Responses via PD-1/PD-L1 Axis. Front. Immunol. 9, 2788 (2018).

124. Jin, X. et al. Identification of shared characteristics in tumor-infiltrating T cells across 15 cancers. Mol. Ther. - Nucleic Acids 32, 189–202 (2023).

125. Na, K. et al. CD81 and CD82 expressing tumor-infiltrating lymphocytes in the NSCLC tumor microenvironment play a crucial role in T-cell activation and cytokine production. Front. Immunol. 15, 1336246 (2024).

126. Tateno, H. et al. Human ZG16p recognizes pathogenic fungi through non-self polyvalent mannose in the digestive system. Glycobiology 22, 210–220 (2012).

127. Zhang, C., Zhao, Z., Liu, H., Yao, S. & Zhao, D. Weighted Gene Co-expression Network Analysis Identified a Novel Thirteen-Gene Signature Associated With Progression, Prognosis, and Immune Microenvironment of Colon Adenocarcinoma Patients. Front. Genet. 12, (2021).

128. Meng, H., Ding, Y., Liu, E., Li, W. & Wang, L. ZG16 regulates PD-L1 expression and promotes local immunity in colon cancer. Transl. Oncol. 14, 101003 (2020).

129. McKenzie, D. R., Comerford, I., Silva-Santos, B. & McColl, S. R. The Emerging Complexity of γδT17 Cells. Front. Immunol. 9, (2018).

130. Mestas, J. & Hughes, C. C. W. Of Mice and Not Men: Differences between Mouse and Human Immunology. J. Immunol. 172, 2731–2738 (2004).

131. Siegers, G. M. et al. Different composition of the human and the mouse γδ T cell receptor explains different phenotypes of CD3γ and CD3δ immunodeficiencies. J. Exp. Med. 204, 2537– 2544 (2007).

132. Qu, G. et al. Comparing Mouse and Human Tissue-Resident γδ T Cells. Front. Immunol. 13, (2022).

133. Brubaker, D. K. et al. An interspecies translation model implicates integrin signaling in infliximab-resistant inflammatory bowel disease. Sci. Signal. 13, eaay3258 (2020).

134. Brubaker, D. K., Proctor, E. A., Haigis, K. M. & Lauffenburger, D. A. Computational translation of genomic responses from experimental model systems to humans. PLoS Comput. Biol. 15, e1006286 (2019).

135. Normand, R. et al. Found In Translation: a machine learning model for mouse-to-human inference. Nat. Methods 15, 1067–1073 (2018).

136. Poussin, C. et al. The species translation challenge-a systems biology perspective on human and rat bronchial epithelial cells. Sci. Data 1, 140009 (2014).

137. Hylander, B. L., Repasky, E. A. & Sexton, S. Using Mice to Model Human Disease: Understanding the Roles of Baseline Housing-Induced and Experimentally Imposed Stresses in Animal Welfare and Experimental Reproducibility. Anim. Open Access J. MDPI 12, 371 (2022).

138. Pelka, K. et al. Spatially organized multicellular immune hubs in human colorectal cancer. Cell 184, 4734–4752.e20 (2021).

139. Khaliq, A. M. et al. Refining colorectal cancer classification and clinical stratification through a single-cell atlas. Genome Biol. 23, 113 (2022).

140. Ji, L. et al. scRNA-seq of colorectal cancer shows regional immune atlas with the function of CD20+ B cells. Cancer Lett. 584, 216664 (2024).

141. Yang, M. et al. Single-cell analysis reveals cellular reprogramming in advanced colon cancer following FOLFOX-bevacizumab treatment. Front. Oncol. 13, 1219642 (2023).

142. Liu, X. et al. Th17 Cells Secrete TWEAK to Trigger Epithelial-Mesenchymal Transition and Promote Colorectal Cancer Liver Metastasis. Cancer Res. 84, 1352–1371 (2024).

143. Hsu, W.-H. et al. Oncogenic KRAS Drives Lipofibrogenesis to Promote Angiogenesis and Colon Cancer Progression. Cancer Discov. 13, 2652–2673 (2023).

144. Guo, W., et al. Resolving the difference between left-sided and right-sided colorectal cancer by single-cell sequencing. JCI Insight 7, e152616 (2022).

145. Zheng, X. et al. Single-cell transcriptomic profiling unravels the adenoma-initiation role of protein tyrosine kinases during colorectal tumorigenesis. Signal Transduct. Target. Ther. 7, 60 (2022).

146. Lenos, K. J. et al. Molecular characterization of colorectal cancer related peritoneal metastatic disease. Nat. Commun. 13, 4443 (2022).

147. Becker, W. R. et al. Single-cell analyses define a continuum of cell state and composition changes in the malignant transformation of polyps to colorectal cancer. Nat. Genet. 54, 985–995 (2022).

148. Zhang, L. et al. Lineage tracking reveals dynamic relationships of T cells in colorectal cancer. Nature 564, 268–272 (2018).

149. Zhang, L. et al. Single-Cell Analyses Inform Mechanisms of Myeloid-Targeted Therapies in Colon Cancer. Cell 181, 442–459.e29 (2020).

150. Che, L.-H. et al. A single-cell atlas of liver metastases of colorectal cancer reveals reprogramming of the tumor microenvironment in response to preoperative chemotherapy. Cell Discov. 7, 80 (2021).

151. Joanito, I. et al. Single-cell and bulk transcriptome sequencing identifies two epithelial tumor cell states and refines the consensus molecular classification of colorectal cancer. Nat. Genet. 54, 963–975 (2022).

152. Stuart, T. et al. Comprehensive Integration of Single-Cell Data. Cell 177, 1888–1902.e21 (2019).

153. Wolf, F. A., Angerer, P. & Theis, F. J. SCANPY: large-scale single-cell gene expression data analysis. Genome Biol. 19, 15 (2018).

154. Hafemeister, C. & Satija, R. Normalization and variance stabilization of single-cell RNA-seq data using regularized negative binomial regression. Genome Biol. 20, 296 (2019).

155. Cao, J. et al. The single-cell transcriptional landscape of mammalian organogenesis. Nature 566, 496–502 (2019).

156. Ran, R., et al. Detailed survey of an in vitro intestinal epithelium model by single-cell transcriptomics. iScience 27, (2024).

